# IGF1R is protective in pneumococcal pneumonia

**DOI:** 10.1101/2024.09.11.612370

**Authors:** Matthias Felten, Luiz-Gustavo Teixeira Alves, Eleftheria Letsiou, Elvira Alfaro Arnedo, Sergio Piñeiro-Hermida, Icíar P. López, Kristina Dietert, Theresa Firsching, Achim D. Gruber, Diana Fatykhova, Andreas Hocke, Tilmann Lingscheid, Jasmin Lienau, Gernot Rohde, Capnetz Study Group, Jose G. Pichel, Martin Witzenrath

**Affiliations:** Charité – Universitätsmedizin Berlin, corporate member of Freie Universität Berlin and Humboldt-Universität zu Berlin, Department of Infectious Diseases, Respiratory Medicine and Critical Care, Berlin, Germany; Berlin Institute for Medical Systems Biology (BIMSB), Max Delbrück Center for Molecular Medicine in the Helmholtz Association (MDC), Berlin, Germany; Division of Pulmonary, Critical Care, Sleep and Allergy, University of Illinois at Chicago, Chicago, IL, USA; Fundación Rioja Salud, Centro de Investigación Biomédica de La Rioja (CIBIR), Lung Cancer and Respiratory Diseases Unit, Logroño, Spain; Instituto de Ciencias de la Vid y del Vino (ICVV), CSIC-UR-Gobierno de La Rioja, Logroño, Spain; Freie Universität Berlin, Department of Veterinary Pathology, Berlin, Germany; Department of Respiratory Medicine, Medical Clinic I, Goethe University Hospital, Frankfurt/Main, Germany; Capnetz Foundation, Hannover, Germany; Biomedical Research in Endstage and Obstructive Lung Disease Hannover (BREATH), Member of the German Center for Lung Research (DZL); Biomedical Research Networking Center in Respiratory Diseases (CIBERES), ISCIII, Madrid, Spain; German Center for Lung Research (DZL), Berlin, Germany

## Abstract

**Background:** *Streptococcus pneumoniae* (*S.pn*) is the most prevalent causal bacterial pathogen in community-acquired pneumonia. Despite appropriate antimicrobial therapy, pneumococcal pneumonia can progress to acute respiratory distress syndrome where actual therapies are mainly supportive, and the discovery of new molecular targets is needed.

**Objective:** To investigate the role of IGF1R (Insulin-like Growth Factor 1 Receptor) in pneumococcal pneumonia.

**Methods:** *Igf1r*-deficient (*UBC-CreERT2; Igf1r^fl/fl^*) and control (*Igf1r^fl/fl^*) mice were infected with 5×10^6^ *S.pn* (PN36) or PBS (sham infected). Mice were sacrificed 48 h after infection. Pulmonary permeability, local inflammatory response, and pulmonary and extra-pulmonary bacterial loads were analyzed. Further, IGF1R protein expression was determined in human lung tissue after *S.pn* infection and IGF1 and IGF1R levels were determined serum of pneumonia patients.

**Results:** In patients and mice infected with *S.pn*, IGF1 signaling was significantly altered. *Igf1r-*deficient mice had significantly increased pulmonary permeability after infection with increased pulmonary inflammatory cytokine levels, while inflammatory cell recruitment was not altered compared to infected *Igf1r^fl/fl^* control animals. Pulmonary bacterial load was significantly higher in *Igf1r*-deficient mice, and histological analysis confirmed increased alveolar edema and necrosis compared to infected *Igf1r^fl/fl^* control and sham-infected mice. *Ex vivo*, *S.pn* caused a decrease in IGF1R protein expression in human lung tissue.

**Conclusion:** Our results demonstrate a significant regulation of IGF1R in ex-vivo infected human lung tissue and in serum of *S.pn* pneumonia patients. Moreover, pneumonia severity was increased in *Igf1r*-deficient mice upon *S.pn* infection compared to *Igf1r^fl/fl^* control mice, suggesting that IGF1R plays a protective role in pneumococcal pneumonia.

## Introduction

Bacterial community-acquired pneumonia (CAP), commonly caused by the bacterial pathogen *Streptococcus pneumoniae* (*S.pn*), is causing millions of deaths worldwide despite effective antibiotic therapy(1, 2). This acute respiratory infection and the associated inflammatory response can lead to increased pulmonary permeability, edema formation and finally to life-threatening acute respiratory distress syndrome (ARDS)(3). Both direct effects of pathogenic factors and an uncontrolled host response can drive pneumonia into acute respiratory failure. The pore forming toxin pneumolysin (PLY) is a major virulence factor of *S.pn*, known to promote alveolar epithelial cell injury, enhance permeability and promote bacteremia(4). An effective host response upon *S.pn* infection is necessary to eradicate the bacteria, however, if not controlled, an excessive local and systemic inflammatory response can further aggravate lung injury and result in lethal sepsis(3).

The insulin-like growth factor 1 receptor (IGF1R) is a widely expressed tyrosine kinase receptor and a central member of the IGF1 axis, that impacts pathways directing cellular growth, differentiation and proliferation (5, 6). The IGF1/IGF1R signaling system is important in lung homeostasis and has been implicated in pulmonary diseases such as asthma, fibrosis, COPD and ARDS (7). Both IGF1- and IGF1R-deficient mice exhibit impaired lung development due to altered alveologenesis and die prematurely (8, 9). Interestingly, our recent studies provide evidence that *Igf1r* plays an important role for the initiation and aggravation of pulmonary inflammation in bleomycin-induced lung injury and lung cancer tumor microenvironment as well as in murine asthma, mediating airway hyperresponsiveness, tissue remodeling and mucus secretion (10–13). On the other hand, Han et al. have reported that conditional deletion of IGF1R in airway epithelial cells led to exacerbated lung inflammation after allergen exposure (14). IGF1 was reported to regulate acute inflammatory lung injury mediated by influenza virus infection, and IGF1R was recently identified as an entry receptor for respiratory syncytial virus and as a novel outcome biomarker in critical COVID-19 patients to predict mortality (15–17). Moreover, studies in ARDS patients provide evidence that IGF1 signaling is involved in profibrotic changes following acute lung injury and even proposed IGF1 inhibition in the treatment of COVID-19-related adult respiratory distress syndrome (18). However, inhibiting IGF1/IGF1R signaling may be a double-edged sword for future therapeutic options, considering possible detrimental effects during lung infections. Actually, Oherle *et al.* have recently reported that IGF1 signaling coordinates concurrent embryonic lung development and mucosal defenses, protecting mice against neonatal respiratory infections (19).

In this study, we employed transgenic mice with an inducible deletion of *Igf1r* activated by tamoxifen (*UBC-CreERT2; Igf1r^fl/fl^* transgenic mice (20) to study the role of IGF1R in pneumococcal pneumonia of adult mice. *Igf1r-*deficient mice showed increased susceptibility to pneumococcal pneumonia with severe barrier disruption as well as increased inflammatory reaction. IGF1R protein expression was down-regulated in human alveolar epithelial cells and human lung tissue exposed to *S.pn*. Moreover, IGF1 and IGF1R plasma levels were found significantly altered in patients with pneumococcal pneumonia. Our results provide new evidence for a protective role of IGF1R in pneumococcal pneumonia and may open the way for future therapeutic approaches targeting the IGF1 axis.

## Methods

For details refer to online supplements.

### Patient serum

CAPnetz provided serum samples of patients with CAP where *S.pn* or influenza virus was identified. The study has been approved by the ethics committee of the Hannover Medical School (registration number 301-2008) and by the local responsible ethics committees of all study centers. Serum levels of IGF1 and soluble IGF1R were quantified by ELISA (Abcam).

### Murine pneumonia model

Animal experiments were reviewed and approved by LAGeSo, Berlin, Germany (Ref. G0300/17) and CIBIR Bioethics Committee Logroño, Spain (Ref. 03/12). Female *Igf1r-*deficient mice and littermate controls (10-12 weeks, 18 to 20 g)(20) were anesthetized with ketamine (80 mg/kg) and xylazine (25 mg/kg) and intranasally inoculated with 5 × 10^6^ colony-forming units (CFUs) of *S. pneumoniae* serotype 3 (PN36, NCTC7978) or 20 μl PBS (sham infection) as described (21, 22).

### Sampling, cell isolation and histology

Mice were anesthetized 48h post-infection (160 mg/kg ketamine and 75 mg/kg xylazine) and exsanguinated prior to harvest. Blood samples were analyzed using a blood-gas analyzer (ABL-800; Radiometer). Bronchoalveolar lavage (BAL) was performed by instillation and drainage of 800 μl PBS. Permeability was assessed by determining total protein concentration (DC protein assay kit, BIO-RAD, USA) in BAL fluid (BALF). Right lung lobes were divided into two parts, homogenized and used for bacterial load determination and leukocyte analysis. Spleen and liver were also homogenized for bacterial load determination. FACS Canto II flow cytometer (BD Bioscience) was used to count and differentiate BAL and lung leukocytes (23). Lung tissue, BALF and blood plasma were stored at −80°C for further analysis. Lung tissue for histological examination was obtained from independent experiments. qPCR was performed with predesigned primers (Qiagen, QuantiTect Primer Assays).

### Bone marrow isolation and reactive oxygen species (ROS) quantification

To isolate cells from bone marrow, femur and tibia were flushed and neutrophils (PMN) enrichment was performed with the EasySep mouse neutrophil enrichment kit (Stem Cell Technologies). PMN ROS production upon stimulation with PMA (100 nM; Thermo Fisher Scientific) or PLY (100 ng/ml) was assessed with a luminol kinetic assay(24). Further, PMNs were incubated with live GFP-tagged *S.pn* (MOI 20, 1 h) and phagocytosis and killing capacity was assessed via flow cytometry.

### *Ex vivo* human lung tissue and *in vitro* human airway epithelial cell model

Human airway epithelial cells (A549 cells, ATCC CCL-185) infected with live D39 wild-type (*S.pn* WT) or D39 pneumolysin-deficient (ΔPly) *S.pn* at MOI 50 for 4h. In separate experiments, A549 were challenged with PLY (100 ng/ml) for 4h. Detailed procedures were described previously (25). Additionally, for infection with *S.pn*, tumor-free human lung tissue was obtained from patients undergoing lung resection. Written informed consent was obtained from all patients. The local ethics committee approved the study (EA2/079/13). Lung organ cultures were inoculated with control or S.pn (1×10^6^ CFU/ml) containing medium for 24h and lungs were processed for western blot analysis with antibodies against IGF1R (Cell Signaling Tech) or β-actin (Sigma-Aldrich).

### Statistical analysis

Statistical analysis was performed using GraphPad Prism 9.00 (USA) software. *P* values <0.05 were considered statistically significant.

## Results

### The IGF1 pathway is regulated in humans and mice during *S.pn* pneumonia

IGF1 is the major ligand of IGF1R and previous studies have shown that IGF1 levels are altered in ARDS (26). Therefore, we measured IGF1 and soluble IGF1R levels in serum samples from patients with pneumonia from the CAPnetz cohort (Figure **1A**). Patients with pneumococcal pneumonia exhibited lower serum levels of IGF1 compared to controls (healthy volunteers). In patients with viral pneumonia (H1N1), IGF1 levels were even slightly, but significantly (p<0.01) lower compared to both controls and pneumococcal pneumonia patients. Importantly, serum levels of IGF1R were significantly elevated in patients with pneumococcal pneumonia as compared to controls (3-fold) and influenza infected patients (2-fold), suggesting a specific regulation of IGF1R signaling in bacterial pneumonia.

**Figure 1.**
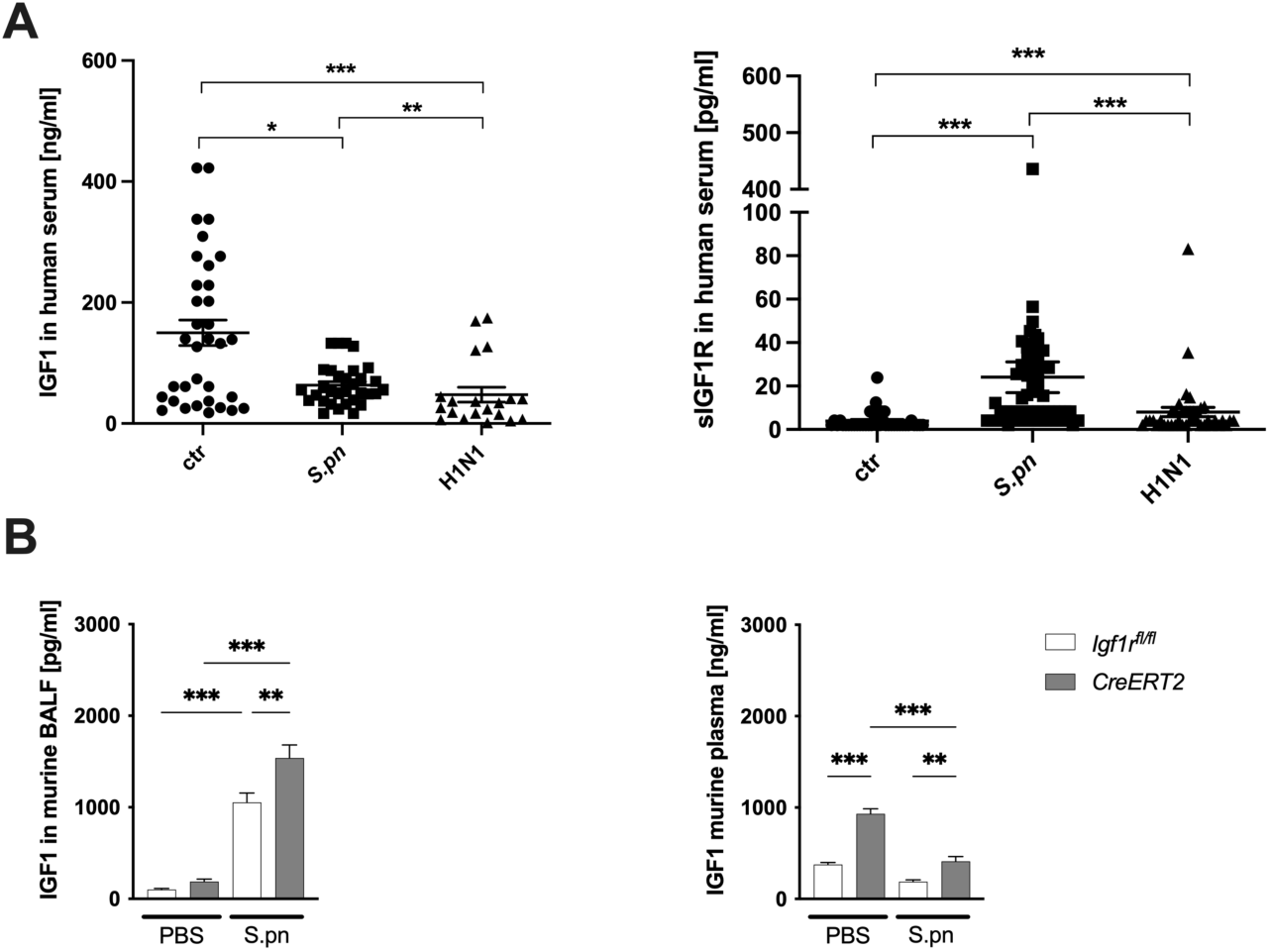
IGF1 blood levels decreased in humans and mice during *S.pn* pneumonia. (**A**) IGF1 (n = 32 for *S.pn*, n = 20 for H1N1 and n = 51 for ctr) and soluble IGF1R (sIGF1R) (n = 62 for *S.pn*, n = 40 for influenza (H1N1) and n = 40 for ctr) levels were determined by ELISA in serum samples from pneumonia patients of the CAPnetz study group and compared to samples from healthy volunteers (ctr). **(B)** *CreERT2* and *Igf1r^fl/fl^*mice were infected with *S.pn* or sham-infected with PBS and sacrificed 48h post infection. IGF1 levels in BALF and plasma were determined by ELISA (n = 5-6 for PBS, n = 8-9 for *S.pn*). Data are median ± SEM. *p < 0.05; ** p < 0.01; ***p < 0.001. (**A** One-way-Anova/ Tukeyߣs multiple comparison test; **B** multiple Mann-Whitney U-test with Bonferroni correction).

To study the role of the IGF1 pathway in bacterial pneumonia, we employed our murine *S.pn* pneumonia model. Previously characterized *Igf1r*-deficient (*CreERT2*) mice and control littermates (*Igf1r^fl/fl^*)(20) were infected with *S.pn* or sham-treated with PBS (control group). Indeed, we observed a significant elevation of IGF1 levels in BAL upon pneumococcal infection in *CreERT2* and *Igf1r^fl/fl^*mice compared to PBS controls. Of note, IGF1 levels were significantly higher in *CreERT2* compared to *Igf1r^fl/fl^* mice after *S.pn* infection (Figure **1B**). In contrast, plasma levels of IGF1 were reduced upon *S.pn* infection. Notably, *CreERT2* mice exhibited increased levels of IGF1 in plasma with respect to PBS or *S.pn*-challenged *Igf1r^fl/fl^* mice. These results suggest a significant regulation of the IGF1 / IGF1R axis during *S.pn.* pneumonia.

### *Igf1r* deficiency in mice enhanced severity of *S.pn* pneumonia

To asses clinical pneumonia severity in IGF1R deficient mice, body weight loss and temperature were monitored every 12 h for a total of 48 h in *CreERT2* and *Igf1r^fl/fl^* animals. In both groups of *S.pn*-infected mice, body weight decreased steadily compared to PBS-treated mice. Specifically, 24 h after infection, *CreERT2* mice experienced greater weight loss compared to *Igf1r^fl/fl^* (p=0.0592), while no differences were observed after 36 - 48 h (Figure **2A**). Mean body temperature in both groups of *S.pn*-infected mice was decreased to 28-30 °C after 12 h compared to PBS-challenged mice (significance not indicated in the graph), and subsequently increased to 32 °C at 48 h. Of note, 12 h after infection *CreERT2 S.pn*-infected mice had a significantly lower body temperature compared to their *Igf1r^fl/fl^*counterparts (Figure **2B**), indicating increased clinical severity during the early phase of *S.pn* pneumonia. To test whether the clinical aggravation was associated with increased pulmonary bacterial burden, we quantified *S.pn* CFUs (colony forming units) in lung homogenates after 48 h and observed that loss of *Igf1r* led to a significant increase in bacterial growth (Figure **2C**).

**Figure 2.**
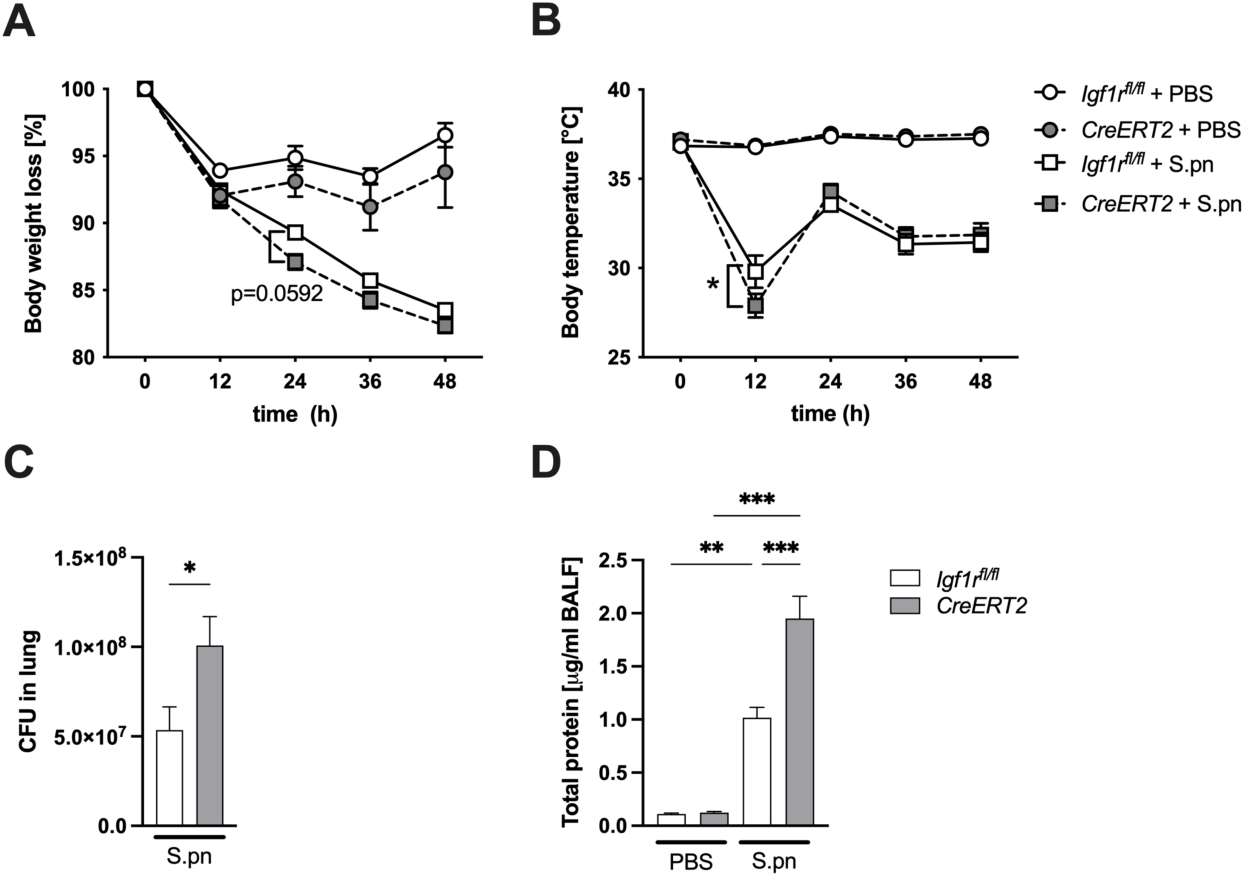
*Igf1r* deficiency in mice enhanced clinical severity of *S.pn* pneumonia in mice. *UBC-Cre-ERT2*; *Igf1r*^fl/fl^ (*CreERT2*) and *Igf1r*^fl/fl^ mice were intranasally infected with *Streptococcus pneumoniae* (*S.pn*) or sham-infected with PBS and sacrificed 48 h post infection. Body weight loss (**A**) and body temperature (**B**) were assessed twice daily until the end point at 48 h in all experimental groups. Bacterial counts (**C**) (colony forming units, CFU) in lung homogenates. Lung permeability (**D**) was assessed by total protein quantification in BAL fluid. Data are mean ± SEM n=5 (PBS), n=8-9 (*S.pn*),*p < 0.05, **p < 0.01, ***p < 0.001 (**A, B** Two-way ANOVA/ Tukey’s multiple comparisons test; **C** Mann-Whitney U-Test; **D**, One-way ANOVA/ Tukey’s multiple comparisons test)

In accordance with disease severity, *S.pn* infection led to lung barrier disruption, as indicated by increased total protein concentration in BALF in respect to PBS-treated mice after 48h. Interestingly, *S.pn*-infected *Igf1r*-deficient mice exhibited significantly higher total protein concentration in BALF compared to *Igf1r^fl/fl^* mice, suggesting a more severe disruption of the alveolar-capillary barrier in these mice (Figure **2D**).

### *Igf1r* deficiency exacerbated pulmonary inflammation and decreased neutrophil influx during *S.pn* pneumonia

Activation the of immune system via inflammatory mediators and rapid recruitment of innate immune cells into the lungs are key features of acute lung injury during bacterial pneumonia. To analyze the effect of *Igf1r* deficiency on the release of inflammatory mediators during *S.pn* pneumonia, we quantified cytokine levels of tumor necrosis factor α (TNFα), monocyte chemoattractant 1 (MCP-1/CCL2), interleukin 1β (IL-1β), interleukin 6 (IL-6), granulocyte-macrophage colony-stimulating factor (GMC-SF), and keratinocyte chemoattractant (KC/CXCL1) in BALF of *CreERT2* and *Igf1r^fl/fl^*mice (Figures **3A**). PBS-treated animals displayed only baseline levels with no significant differences between groups (Figure **S1**). However, *S.pn* infection significantly increased cytokine levels in both mouse genotypes. Interestingly, although TNFα and IL-6 did not show differences between genotypes, BALF levels of MCP-1/CCL2, IL-1β, GMC-SF, and KC/CXCL1 levels were significantly higher in the *CreERT2* compared to *Igf1r^fl/fl^* mice upon *S.pn* pneumonia (Figures **3A**). These results are in line with the increased clinical severity observed in *S.pn*-challenged *Igf1r-*deficient animals. To verify if the increased pulmonary cytokine release led to an increased immune cell recruitment, we performed a flow cytometry analysis of BAL and lung tissue. While we observed increased neutrophil (PMN, polymorphonuclear leukocytes) and inflammatory monocyte numbers in BAL upon *S.pn* infection, their counts did not show significant differences between *S.pn*-infected *Igf1r^fl/fl^* and *CreERT2* (Figure **3B**, top). However, we detected significantly lower PMN neutrophil counts in lung homogenates of *Igf1r-*deficient mice compared to *Igf1r^fl/fl^* mice upon *S.pn* infection, while alveolar macrophage and inflammatory monocyte counts exhibited no difference between experimental groups (Figure **3B**, bottom). Taken together our findings suggest that *Igf1r*-deficiency exacerbates pneumonia-induced lung injury and inflammation, which could be associated with an impaired recruitment of PMN (neutrophils).

**Figure 3.**
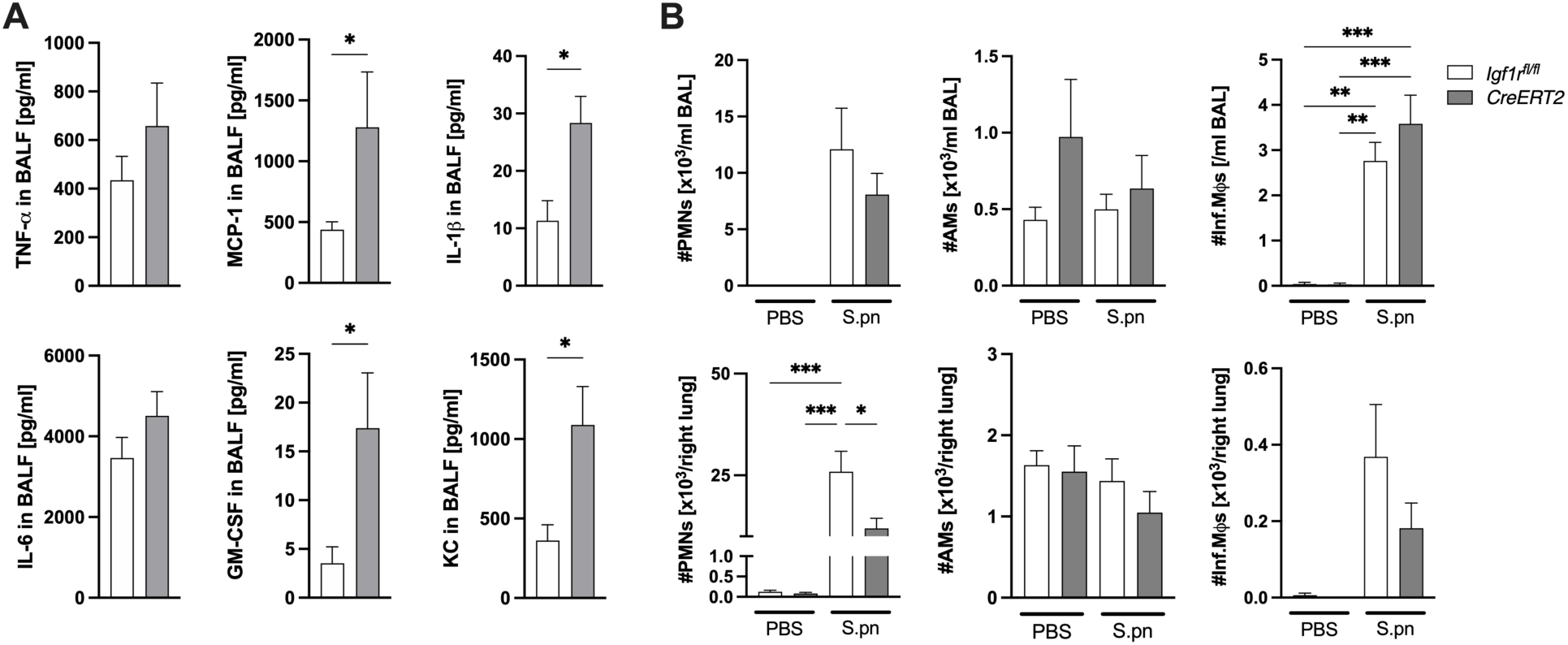
*Igf1r* deficiency exacerbates pulmonary inflammation and decreases neutrophil influx in BAL during *S.pn* pneumonia. *CreERT2* and *Igf1r*^fl/fl^ mice were intranasally infected with *S.pn* or sham-infected with PBS and sacrificed 48 h post infection. (**A**) Inflammatory cytokines in *S.pn* infected animals were quantified via multiplex ELISA in BAL fluid (BALF). (**B**) Quantifications of polymorphonuclear neutrophils (#PMNs), alveolar macrophages (#AMs) and inflammatory monocytes (#inf.MՓs) were assessed by flow cytometry in BAL (top) and lung tissue (botton). Data are mean ± SEM. n = 5 (PBS), n = 8-9 (*S.pn*), *p < 0.05, **p < 0.01, **p < 0.001 (**A** Mann-Whitney U-Test**; B** One-way ANOVA/ Tukey’s multiple comparisons).

### *Igf1r* deficiency led to increased histological signs of lung injury and reduced neutrophil survival upon *S.pn* infection

To corroborate our findings, we histologically evaluated the lungs of *CreERT2* and *Igf1r^fl/fl^* mice 48 h after *S.pn* infection (Figure **4**). Histopathologic examination revealed increased signs of alveolar edema, tissue necrosis and vasculitis in *CreERT2* mice. Interestingly, morphologic analysis of PMNs in *Igf1r^fl/fl^*mice unveiled predominantly healthy PMNs, whereas in *Igf1r*-deficient mice, fragmented necrotic PMNs were prevalently found evenly distributed in the infected lung areas (Figure **4A**). Semiquantitative analysis confirmed a significantly higher score for necrosis and steatitis, as well as tendence to show increased alveolar edema severity (p = 0.0794) in *CreERT2* compared to *Igf1r^fl/fl^* mice (Figure **4B**).

**Figure 4.**
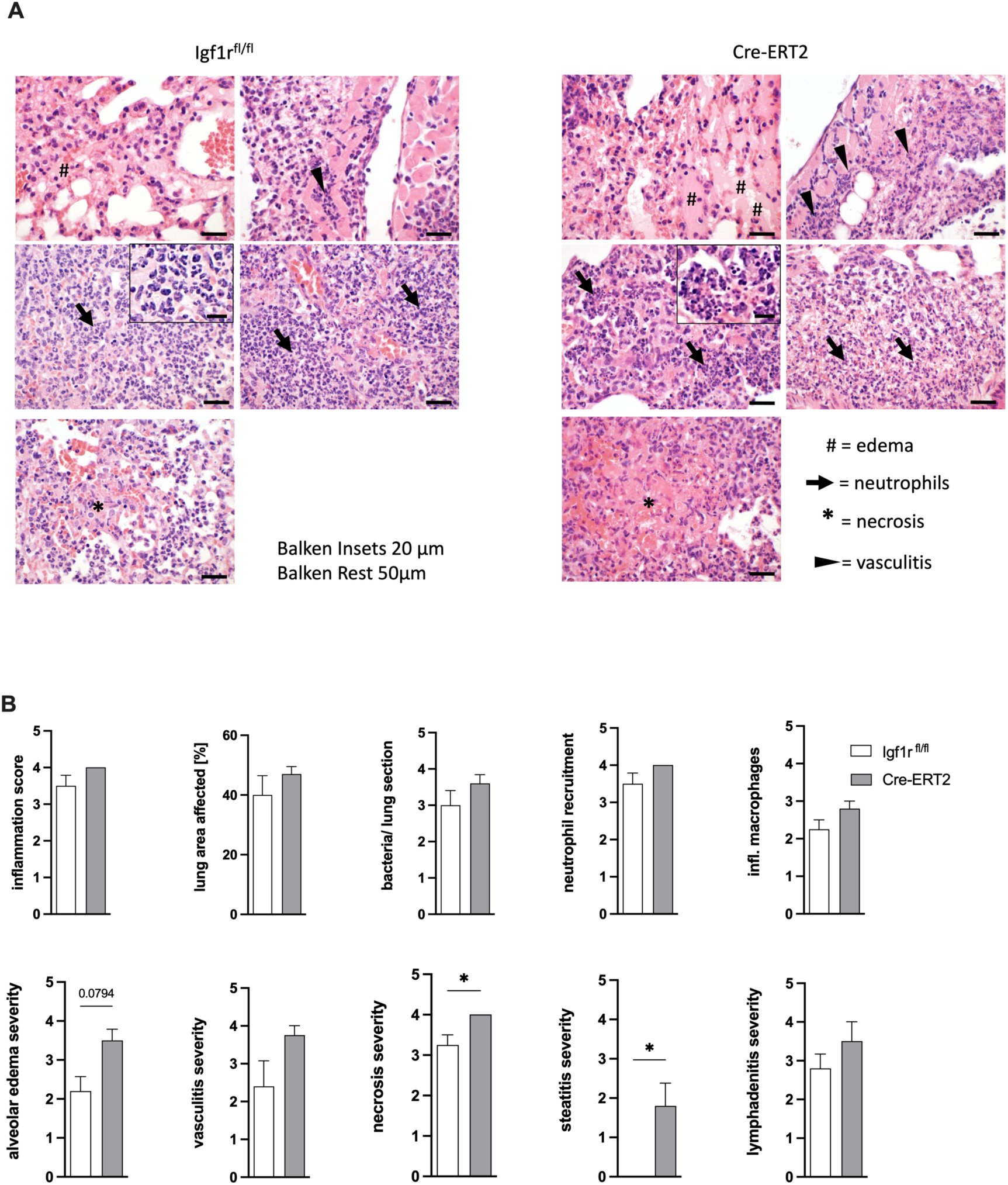
*Igf1r* deficiency led to increased alveolar edema, necrosis and steatitis upon *S.pn* infection. *CreERT2* and *Igf1r^fl/fl^* mice were intranasally infected with *S.pn* and sacrificed 48 h post infection. Paraffin-embedded lung sections were stained with hematoxylin and eosin. (**A**) Representative histologic images show that *CreERT2* mice developed more severe alveolar edema (#), increased signs of vasculitis (►) and predominantly fragmented necrotic neutrophils (→) to viable neutrophils compared with *Igf1r^fl/fl^* animals. Enhanced signs of tissue necrosis (*) were also observed in *Igfr1*-deficient animals. Scale bars: 50 µm, 20 µm in insets. (**B**) Quantification of histologic signs of lung injury: total lung inflammation score, percentage of lung area affected by *S.pn* pneumonia, bacteria per lung section, neutrophil infiltration, inflammatory macrophages, alveolar edema, vasculitis, necrosis, steatitis and lymphadenitis. All scoring parameters were rated as 0: non-existent, 1: minimal, 2: mild, 3: moderate, 4: severe. Data are mean ± SEM. n = 4 per condition. *p < 0.05 (Mann-Whitney U-Test).

### *Igf1r* deficiency increased systemic inflammation and decreased blood neutrophil counts upon S.pn infection

Severe pneumonia and associated lung barrier disruption can lead to a systemic dissemination of bacteria, enhance systemic inflammation and drive extrapulmonary organ failure. Blood, liver and spleen CFU did not show significant differences between *S.pn*-infected *CreERT2* and *Igf1r^fl/fl^* mice (Figure **5A**). However, we observed a tendency towards increased plasma levels of aspartate transaminase (AST, used as a surrogate marker for liver function) in the *S.pn* infected *CreERT2* mice compared to *Igf1r^fl/fl^* controls (p = 0.069) indicating moderate extrapulmonary organ injury (Figure **5B**). Yet, plasma levels of urea (marker for kidney function) did not exhibit significant differences between experimental groups (**Figure 5B**). To characterize extrapulmonary inflammation we quantified cytokine levels in plasma and performed a flow cytometry analysis of inflammatory cells (Figure **5C, D**). Plasma levels of TNF-α, IL-1β, IL-6, GM-CSF and KC did not significantly alter upon *S.pn* infection. Specifically, only MCP-1 levels were significantly incremented in *S.pn*-infected *CreERT2* compared to *Igf1r^fl/fl^* mice (**Figure 5C**). PBS-treated animals displayed baseline levels for these markers with no significant differences between groups (Figure **S2**). In line with our observation of PMN decreased neutrophil counts in lung tissue, PMN counts in blood decreased by 50% in *S.pn*-infected *Igf1r-*deficient mice compared to *Igf1r^fl/fl^.* On the other hand, PMN counts in liver, as well as inflammatory monocytes in both blood and liver were not statistically significant between experimental groups (**Figure 5D**).

**Figure 5.**
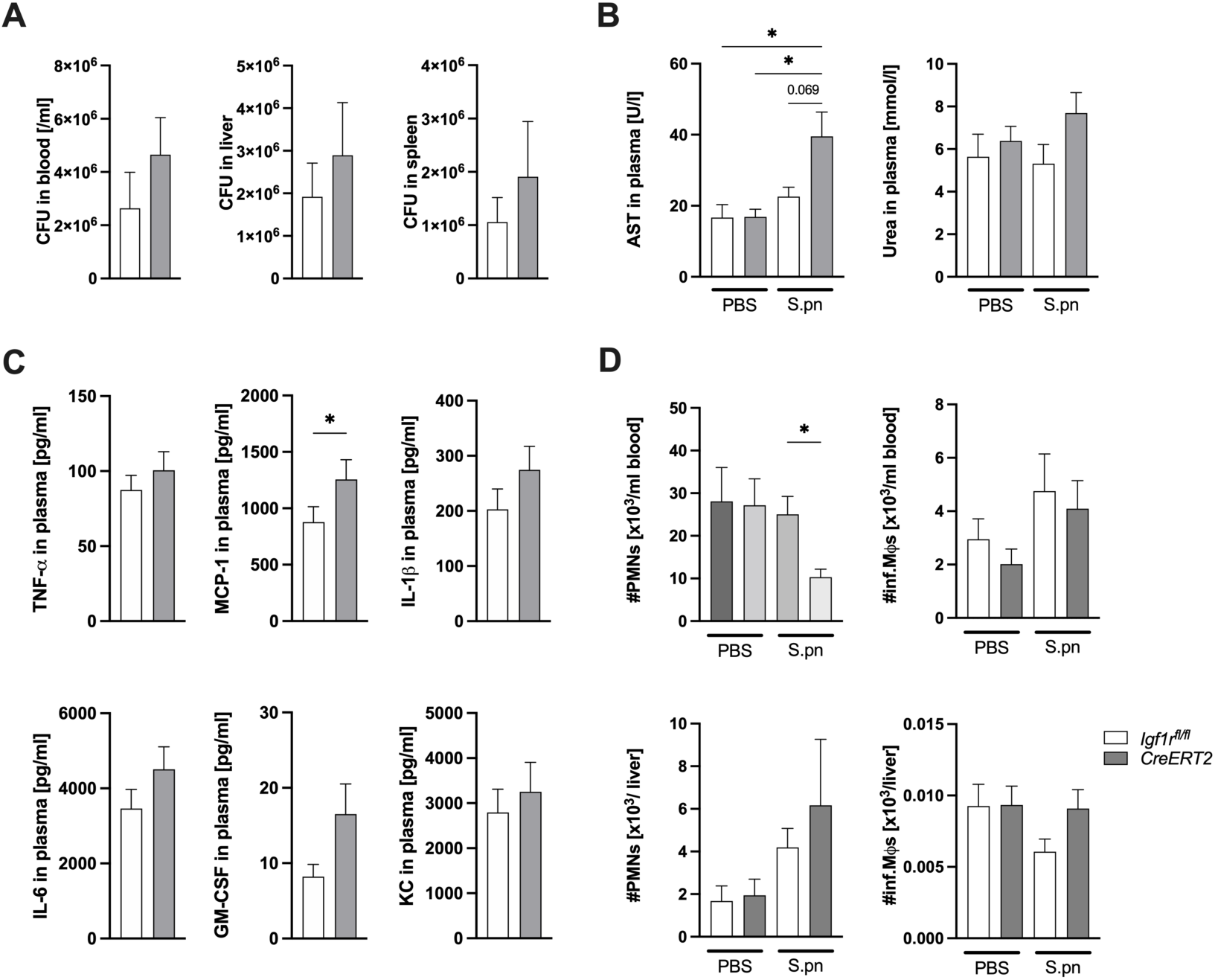
*Igf1r* deficiency increased systemic inflammation and decreased blood neutrophil counts upon *S.pn* infection. *CreERT2* and *Igf1r*^fl/fl^ mice were intranasally infected with *S.pn or* sham infected with PBS (controls) and sacrificed 48 h post infection. (**A**) Extrapulmonary bacterial dissemination in infected animals was quantified via bacterial counts (colony forming units, CFU) in blood, liver and spleen homogenates of infected animals. (**B**) Clinical parameters: AST (aspartate transaminase, used as a surrogate marker for liver function) and urea (marker for kidney function) were measured in plasma. (**C**) Inflammatory cytokines in infected animals were quantified by multiplex ELISA in plasma. (**D**) Quantification of polymorphonuclear neutrophils (PMNs), and inflammatory monocytes (Infl.Mϕs) were assessed by flow cytometry, and numbers (#) of polymorphonuclear neutrophils (PMN), and inflammatory monocytes (infl.M) were calculated. Data are mean ± SEM. n = 5 (PBS), n = 8-9 (*S.pn*); *p < 0.05 (**A** and **C** Mann-Whitney U-Test, **B** and **D** One-way ANOVA/ Tukey’s multiple comparison test).

### *Igf1r* deficiency decreased neutrophil activation without affecting antibacterial function

Taking into account the observed decrease in PMN counts upon *S.pn* infection in lung and blood of *CreERT2* mice, we next aimed to investigate whether *Igf1r* deficiency affects PMN function *in vivo and in vitro*. First, we assessed the expression of the PMN activation marker CD11b on blood PMNs by flow cytometry. CD11b surface expression was significantly lower on PMNs of PBS-treated *CreERT2* compared to *Igf1r^fl/fl^* mice, while upon *S.pn* infection the difference was dampened showing high levels in both groups (Figure **6A**). Next, we investigated whether *Igf1r* deficiency affects the phagocytic and killing capacity of PMNs. To address this, PMNs were isolated from bone marrow of *CreERT2* and *Igf1r^fl/fl^* mice and infected with GFP-tagged *S.pn*. We did not observe any difference between PMNs from *Igf1r^fl/fl^* and *CreERT2* in their ability to phagocytose bacteria (Figure **6B**). Interestingly, we observed a significantly lower bacterial killing capacity of PMNs isolated from *CreERT2* mice compared to *Igf1r^fl/fl^* mice (Figure **6B**). However, neutrophilic ROS production upon stimulation with PMA (phorbol myristate acetate) or pneumococcal exotoxin pneumolysin (PLY) did not differ between the experimental groups (Figure **6C**). Since activation of PMNs is required for adequate bacterial clearance, these data suggest that *Igf1r*-deficient PMNs might have an impaired activation, while ROS production and phagocytic capacity remain unaffected.

**Figure 6.**
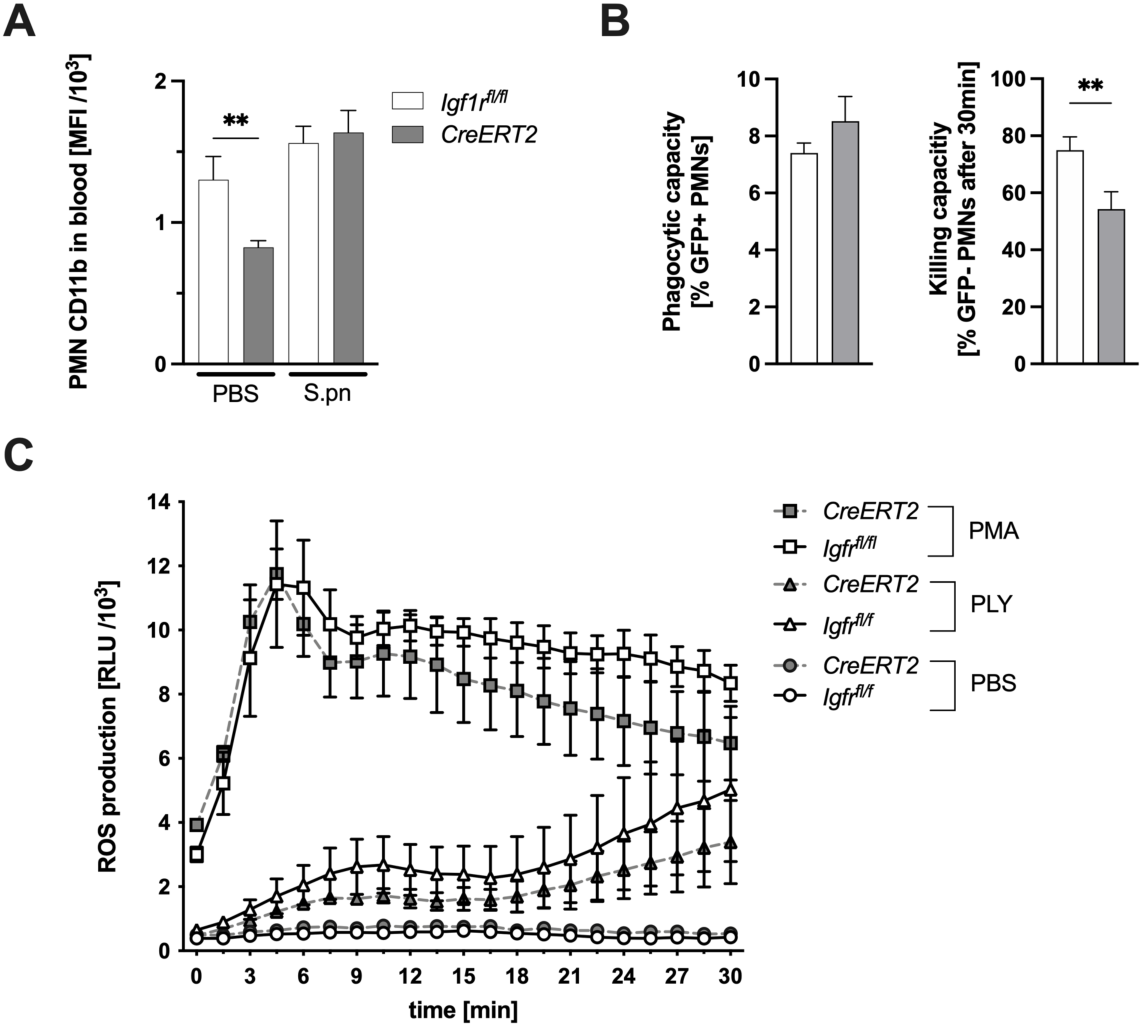
*Igf1r* deficiency impaired neutrophil activation without affecting antibacterial function. *CreERT2* and *Igf1r^fl/fl^* mice were intranasally infected with *S.pn.* or sham-infected with PBS (controls) and sacrificed 48 h post infection. (**A**) Neutrophil activation was measured via CD11b surface expression on blood polymorphonuclear neutrophils (PMNs) by flow cytometry. For functional essays (**B**-**C**), PMNs were isolated from bone marrow cell suspensions from *CreERT2* and *Igf1r^fl/fl^* mice. (**B**) PMNs were incubated with live GFP-tagged *S.pn* (MOI 20, 1 h). Neutrophil phagocytosis and killing capacity was assessed by flow cytometry. (**C**) ROS production after pneumolysin (PLY) or phorbol myristate acetate (PMA) stimulation was monitored over time with fluorophore luminol in a luminescence reader (RLU – *relative light units*). Data are mean ± SEM. n = 3; **p < 0,01 (**A** One-way ANOVA/ Tukey’s multiple comparisons test, **B** Mann-Whitney U-Test and **C** Two-way ANOVA with mixed-effects model/Geissner-Greenhouse correction, Tukey’s multiple comparisons test).

### *S.pn* reduced IGF1R expression in human and murine lung tissue and alveolar epithelial A549 cells

Next, we aimed to determine whether *S.pn* affects IGF1R expression in lung tissue and cells. First, we quantified *Igf1r* mRNA expression levels in mouse lung tissue after *S.pn* infection in *Igf1r^fl/fl^*mice. We found a significant decrease in *Igf1r* mRNA expression compared to PBS controls (Figure **7A**). To verify the translational relevance of our results we employed an *ex vivo* model of human lung tissue infection (27). We observed a significant decrease in IGF1R protein expression after *S.pn* stimulation (Figure **7B**). Consistent with this observation, IGF1R expression was also reduced in *S.pn*-infected A549 cells (Figure **7C**). We have previously demonstrated that *S.pn* mediates its injurious effects by releasing PLY (21). To investigate whether PLY is involved in the regulation of IGF1R expression, we stimulated A549 cells with ΔPLY *S.pn,* a strain that does not express PLY. Interestingly, we observed no difference in IGF1R protein expression in ΔPLY *S.pn*-infected cells compared to control (PBS), suggesting a role of PLY in IGF1R regulation (Figure **7C**). To confirm this finding, we performed additional experiments to quantify the direct effect of PLY on A549 cells and detected a moderate but significant downregulation of IGF1R expression after PLY challenge compared to PBS control (Figure **7D**), suggesting a relevant regulation of IGF1R signaling in pneumococcal pneumonia.

**Figure 7.**
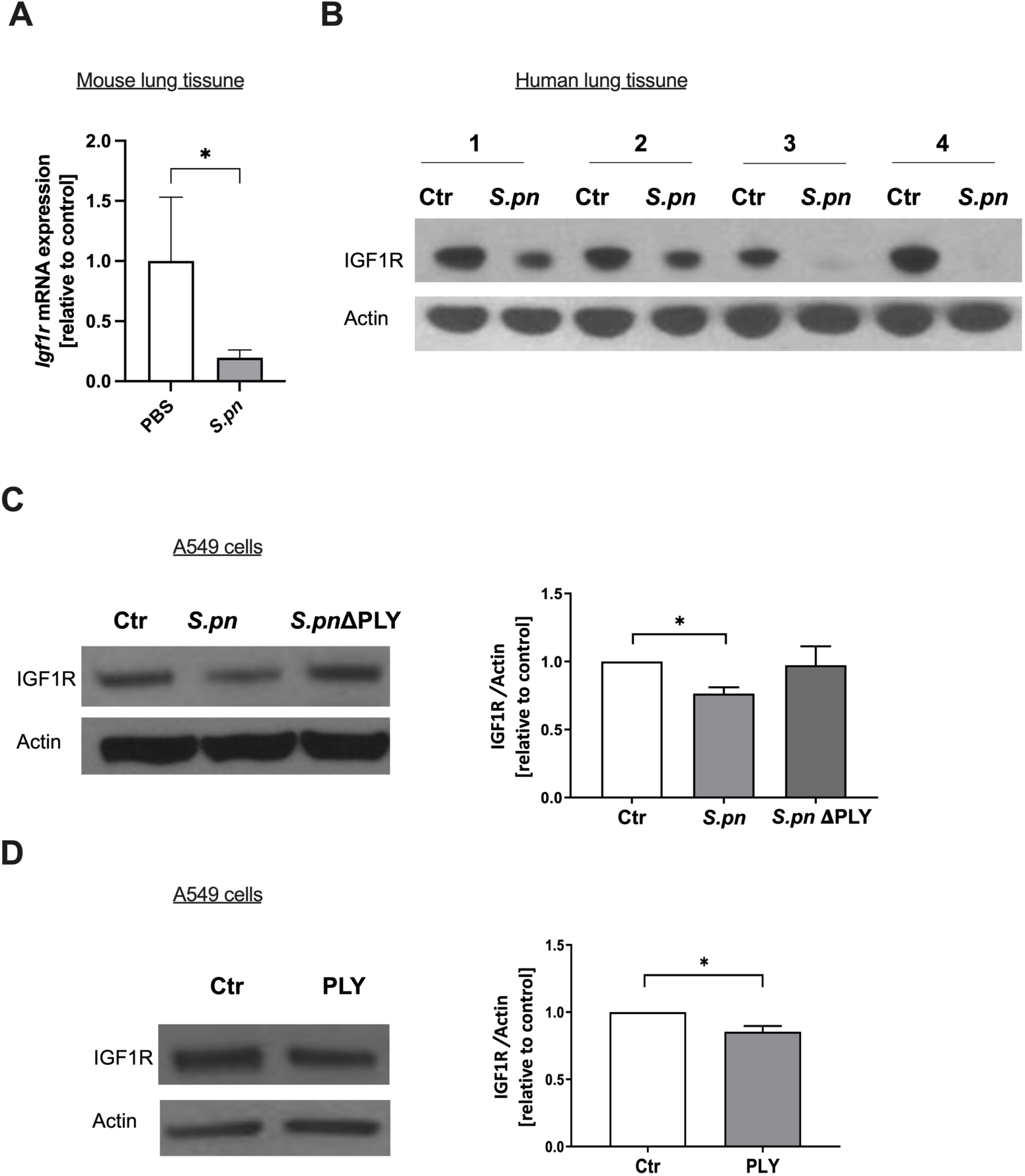
S.pn reduced IGF1R expression in human lung tissue and alveolar epithelial A549 cells. **(A)** *Igf1r^fl/fl^* mice were intranasally infected with *S.pn* PN or sham-infected with PBS and sacrificed 48 h post infection. Gene expression of *Igf1r* was determined by qPCR in lung tissue homogenates (n = 5). (**B**) Human lung specimens (pieces from lung tissue from the same patient) were treated with *S.pn* (MOI 50, 4 hours) by quadruplicate, and IGF1R protein expression was determined by western blotting in lung lysates. (**C-D**) Representative blots and densitometric analysis of IGF1R expression in human alveolar epithelial cells (A549) treated with PBS (Ctr) or stimulated with *S.pn* or a pneumolysin-deficient (ΔPLY) *S.pn* strain (MOI 50, 4 hours) (**C**), or with pneumolysin (PLY; 100 ng/ml, 4 hours) (**D**). Data are mean ± SEM. n = 4. *p < 0.05. (**A** and **C**; Kruskal-Wallis/ Dunn’s post hoc test, **D**; Welch’s t-test).

## Discussion

In this study, we reveal that IGF1R may have a protective role in pneumococcal lung infection. IGF1R deficiency in *S.pn*-infected mice led to aggravated pneumonia with increased bacterial load and lung barrier disruption, reduced blood PMN counts as well as increased inflammatory cytokine production. Moreover, IGF1R protein and gene expression was shown to be downregulated in mouse lungs and *ex vivo* in human lung tissue upon *S.pn* infection, while IGF1 levels increased in BALF of infected mice. On the contrary, IGF1 serum levels decreased during *S.pn* pneumonia in humans and mice.

Disruption of the alveolar capillary barrier is a hallmark of pneumonia-induced acute lung injury (28) and pneumococcal exotoxin PLY is a well-known crucial factor for lung barrier disruption (4). Notably, IGF1 signaling was recently suggested to be involved in the regulation of junction proteins in endothelial cells via IGF1R, protecting endothelial barrier permeability (29). To study the role of IGF1R in pneumonia, we employed a well-established murine model of pneumococcal pneumonia (21, 22). In our study we did not observe differences in permeability in uninfected mice with or without *Igf1r*-deletion, in accordance with our previous results in these mice (10, 30). Upon *S.pn*-infection, *CreERT2* mice exhibited a notably higher lung permeability and more severe histological signs of lung injury and alveolar edema when compared with their *Igf1r^fl/^*^fl^ counterparts. Results shown here suggest that either PLY produced by the increased amounts of *S.pn*, the *Igf1r* deficiency itself in pneumonia conditions or both contribute to the barrier disruption observed in *S.pn*-infected *CreERT2* mice.

To fight infections, a set of immunological responses of critical importance is needed. Pathogen recognition followed by inflammatory mediator release leads to activation of the host innate immune system, and predominantly macrophages and PMNs are recruited to eradicate bacteria (31). Although this inflammatory response is necessary to clear pathogens, excessive inflammatory responses can aggravate lung injury and barrier failure (31, 32). In line with this paradigm, we found increased levels of MCP-1, IL-1ß and GM-CSF in the BAL of *Igf1r*-deficient compared to *Igf1r^fl/fl^*-competent mice after *S.pn* infection. Increased pulmonary MCP-1 levels were found to strongly correlate with bacterial loads in mice (33) and IL-1β was shown to induce endothelial permeability and aggravate lung injury (34, 35). In contrast to our results in pneumococcal pneumonia, inhibition of IGF1 signaling during influenza infection led to a decrease in IL1-ß, TNF-α and IL6 levels, while IGF1 treatment led to increased inflammation (15). Further, proinflammatory pathologies, including acute lung injury evoked by bleomycin, allergic airway inflammation induced by house dust mite (HDM), and lung cancer in a metastasis model were counteracted by *Igf1r*-deficency in *CreERT2* mice (10–12, 30). Additionally, the HDM-induced allergic phenotype was pharmacologically ameliorated in mice treated with an IGF1R inhibitor (13).

The seemingly different findings in pneumococcal pneumonia as compared to other inflammation models made us hypothesize that *Igf1r* deficiency may be associated with impaired innate immune responses to bacterial challenge, leading to increased bacterial growth, and resulting in lung inflammation and injury. Neutrophil activation and recruitment are some of the most crucial components of the initial innate immune response against bacterial infection (36, 37). Preceding studies provided evidence that IGF1 signaling is involved in neutrophil function and may also increase PMN survival and delay apoptosis (38). Accordingly, we observed reduced lung and blood neutrophil numbers in *CreERT2* mice upon *S.pn* infection and decreased bacterial killing capacity *ex vivo*, suggesting an impairment of neutrophil activation due to impaired IGF1R signaling. In the same line, we have previously observed reduced numbers of neutrophils in lungs and BALs of *CreERT2* mice after either an acute bleomycin-induced acute lung injury or a HDM-activated allergy challenge (10, 12). The fact that histological analysis of infected lungs revealed a higher number of apoptotic/necrotic neutrophils in *CreERT2* mice compared to *Igf1r^fl/fl^*controls, may indicate a potential role of IGF1R signaling as antiapoptotic and pro-survival for neutrophils in pneumonia. Interestingly, we observed increased GM-CSF levels in lung tissue of IGF1R deficient mice during *S.pn* infection, a chemotactic signal for neutrophil recruitment, protective in pneumococcal pneumonia (39). In this context GM-CSF secretion may be also interpreted as compensatory mechanism, as it is known to stimulate neutrophil recruitment and granulopoiesis (40). Taking into account our previous studies, where IGF1R has been shown to play a beneficial role in acute and chronic lung inflammation (10–13, 30), these differences compared to the harmful effect in pneumonia are intriguing. It is tempting to speculate, that the observed impairment had detrimental effects in our bacterial infection model, while a dampening of the PMN response led to a better clinical outcome during sterile(10) or viral inflammation (15).

Although the IGF1R ligands IGF1 and IGF2 are primarily produced by the liver (41), both ligands and IGF1R are expressed in the mouse lung, mainly in mesenchymal and epithelial cells (42, 43). Signaling via IGF1R has been shown to enhance cell survival and inhibit lung fibroblast apoptosis (44) and IGF1R deficiency was reported to be protective in lung fibrosis models (10). In the present study, IGF1R protein levels were found decreased in human lung epithelial cells after stimulation with PLY or PLY-producing bacteria, in line with our observations of decreased *Igf1r* gene expression in murine lung tissue upon *S.pn* infection. To date, little is known about the origin and function of soluble IGF1R, which was previously detected in serum and serum-derived exosomes of lung cancer and myocardial infarction patients (30, 45). For the first time, we detected increased serum levels of soluble IGF1R in pneumonia, specifically in patients with *S.pn* pneumonia but not Influenza.

Clinical data on the role of the IGF1-IGF1R system in acute respiratory infections is still controversial. Tejera *et al.* found a significant correlation between high IGF1 serum levels and pneumonia severity in community acquired pneumonia on admittance (46). In contrast, clinical studies in ARDS patients (26) and experimental studies in murine sepsis (47) show that low IGF1 serum levels were associated with poor outcome. Accordingly, we detected decreased serum levels of IGF1 in patients with *S.pn* or Influenza pneumonia compared to healthy controls. It is necessary to consider the dynamic changes in different disease stages, ranging from high IGF1 concentrations in serum and low concentrations in BALF within the first 24h of ALI/ ARDS, that subsequently switch to high alveolar and low serum levels of IGF1 at later stages of lung injury (48). These observations suggest that high IGF1 concentrations in BALF and its decreased serum levels in *CreERT2* mice 48 h after infection may point to state of transition between acute pulmonary inflammation and beginning of sepsis.

Numerous reports in the last decades of IGF research suggest multiple health benefits of inhibiting IGF1R-mediated signaling to counteract, for example, inflammation, tissue fibrosis, cellular senescence, ageing, tumor growth, and others (49–54) or specifically in the lung, to treat asthma, allergy, COPD, lung fibrosis, or lung cancer (7, 55–59). In contrast, this pan-therapeutic approach could have highly malicious effects on early life in the organism, including impairment of DNA repair, cell proliferation and tissue regeneration (60, 61), as well as dampening the host inflammatory defense against pneumonia, as reported here. Therefore, a better understanding of cell- and disease specific regulations of IGF1R signaling appears to be critical for future development of therapeutic application in order to maintain the organismal homeostasis.

Interestingly, while Oherle *et al.* have identified IGF1 as potential mediator of ILC mediated mucosal defenses, which could additionally affect pneumonia (19), recent studies found evidence that enhanced IGF1 production by alveolar macrophages suppressed endogenous inflammatory signals of alveolar epithelial cells mitigating lung injury (62). Accordingly, we found evidence for *S.pn* specific regulation of IGF1R expression in alveolar epithelial cells, but further experimental studies and clinical trials are lacking in order to verify these results and to identify IGF system signaling components that affect cell-specific interactions in acute pneumonia.

In conclusion, our data suggest that IGF1R signaling is significantly decreased in the context of pneumococcal pneumonia and plays an important role in determining the severity of infection *in vivo*. Moreover, we found significant downregulation of IGF1R expression in human lung tissue infected with *S.pn* and increased serum levels of soluble IGF1R in patients with pneumococcal pneumonia. Based on these results, we propose IGF1R regulation at the pulmonary level as key point in pneumococci-induced inflammation and barrier failure in pneumonia. Further investigations are needed to characterize the possible detrimental collateral effects of IGF1R inhibition during bacterial pneumonia.

## Authors’ contributions

MF, GT and EL performed in vivo and in vitro experiments, analyzed data and drafted the manuscript. IPL, EA-A, SP-H and JGP generated and provided experimental mice. EA-A and SP-H performed in vitro experiments. MF, JGP and MW planned and supervised the study, interpreted the data and revised the manuscript. JL performed statistical analysis and revised the manuscript. ADG, KD and TCB performed histological examination and revised the manuscript. TL processed clinical samples provided be GR and the Capnetz Study group.

## Acknowledgements

This work was supported by grants from the Deutsche Forschungsgemeinschaft (DFG, German Research Foundation), SFB-TR84 sub-projects A02, C06, C09 and Z01b, SFB 1449 (431232613) sub-projects B1 and B2, and by the German Ministry of Education and Research (BMBF) in the framework of CAPSyS (01ZX1304B, 01ZX1604B), SYMPATH (01ZX1906A), PROVID (01KI20160A) Phage4Cure (16GW0141), and MAPVAP (16GW0247).

The investigators of this scientific work acknowledge CAPNETZ STIFTUNG and the competence network CAPNETZ for project support with regard to using biomaterials and clinical data. CAPNETZ is a multidisciplinary approach to better understand and treat patients with CAP. The network has been made possible by the contributions of many investigators. CAPNETZ is supported by the German Center for Lung Research (DZL) since 2013, and was supported by the German Ministry of education and Research (BMBF) 2001-2011.

## Supplementary Figures

**Figure S1.**
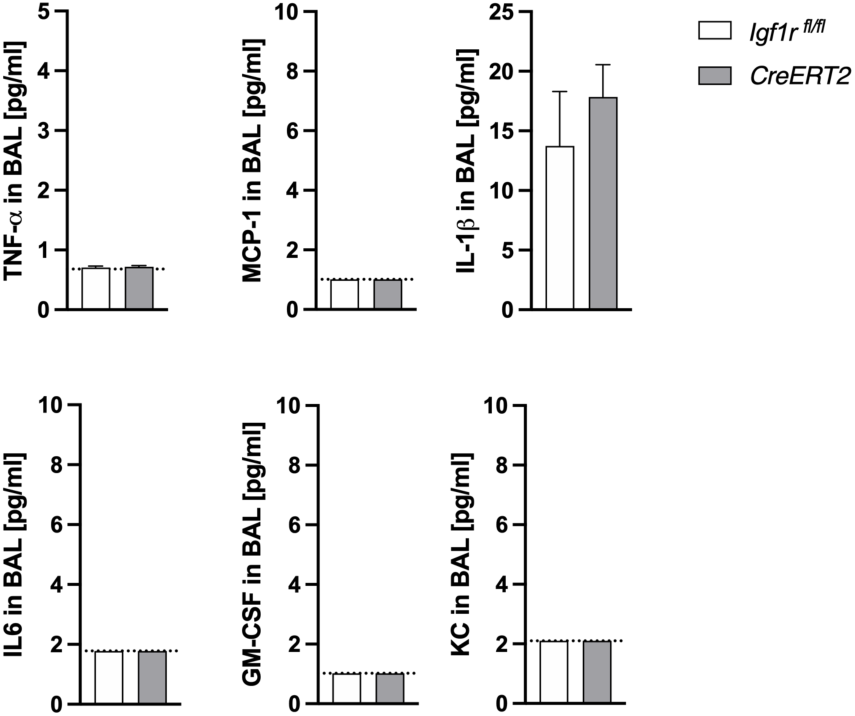
*Igf1r* deficiency does not affect basal levels pulmonary of inflammatory cytokine. *CreERT2* and *Igf1r^fl/fl^* mice were sham-infected with PBS and sacrificed 48h post infection. Inflammatory cytokines were quantified by multiplex ELISA in BAL fluid (BALF). Data are mean ±SEM. n = 5, *p < 0.05 (Mann-Whitney U-Test).

**Figure S2.**
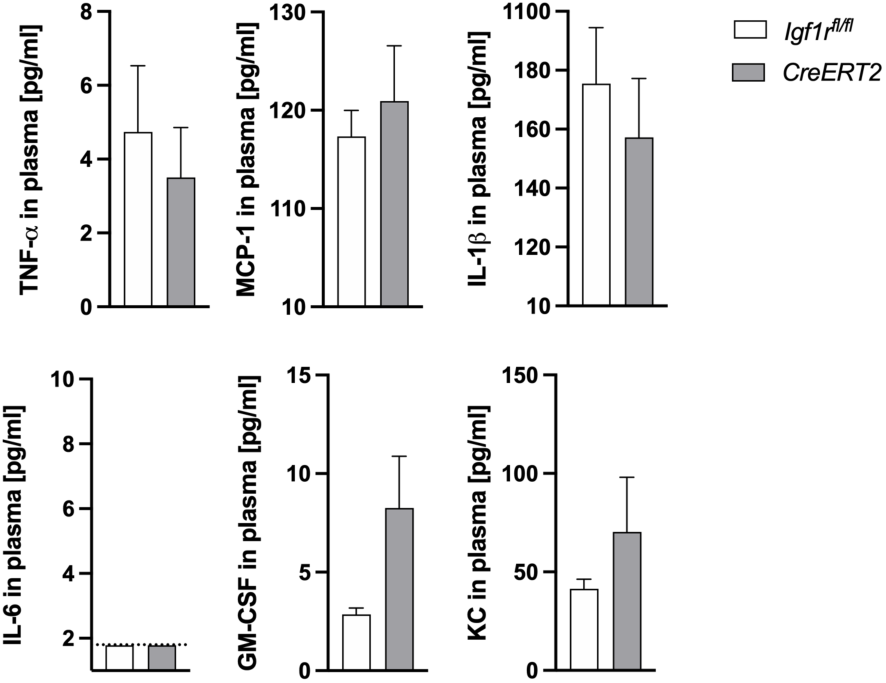
*Igf1r* deficiency does not affect basal plasma levels of inflammatory cytokines. *CreERT2* and *Igf1r^fl/fl^*) were sham-infected with PBS and sacrificed 48h p.i. Inflammatory cytokine levels were quantified by multiplex ELISA in plasma. Data are mean ± SEM. n = 5. (Mann-Withney U-Test).

