## Supplementary material for "IGF1R is protective in pneumococcal pneumonia": online supplements

**Supplementary Materials and methods**

### Animals

All animal studies were approved by the institutional and local governmental authorities at the Charité–Universitätsmedizin Berlin, the Berlin State Office for Health and Social Affairs Berlin, Germany (Ref. G0300/17) and CIBIR Bioethics Committee Logroño, Spain (Ref. 03/12). Experimental procedures were carried out in accordance with the European Directive 2010/63/EU and complied with the Federation of European Laboratory Animal Science Associations (FELASA) guidelines. Experiments were performed using *Igf1r*-deficient mice and littermate controls (10 to 12 weeks old, weighing 18 to 20 g). Animal breeding, tamoxifen induction and maintenance under specific pathogen-free (SPF) conditions were carried out at the CIBIR animal facility). For acclimation in the Berlin animal facility, mice were housed in SPF cages (water and autoclaved food a*d libitum*; 12 h light/dark cycle, 20-23 °C and 40-60 % humidity) for 8-10 days. All handling and preparation procedures were performed under sterile bench conditions.

For experimental purposes, *UBC-Cre-ERT2;Igf1r*$^{fl/fl}$ mice were crossed with *Igf1r*$^{fl/fl}$ mice to directly generate descendants in equal proportions in the same litter. Tamoxifen (TMX) was administered daily for five consecutive days to four-week-old mice of both genotypes, to induce a postnatal *Igf1r* gene conditional deletion in *UBC-Cre-ERT2;Igf1r*$^{fl/fl}$ mice, generating *Igf1r*-deficient mice (now on called *CreERT2*)[15].

### Murine pneumonia model

Mice were anesthetized with ketamine (80 mg/kg) and xylazine (25 mg/kg) and intranasally inoculated with $5 \times 10^6$ colony-forming units (CFUs) of *S. pneumoniae* serotype 3 (PN36, NCTC7978) or 20 µl PBS (sham infection) as previously described

(1, 2). Body weight and temperature were documented twice daily. 48 h post-infection (p.i.), mice were anesthetized (160 mg/kg ketamine and 75 mg/kg xylazine) and exsanguinated prior to harvest. Blood was collected in EDTA tubes and used for hemogram (measured with Scil Vet abc, Scil Animal Care) and bacterial load determination. The remaining blood was centrifuged, and plasma was stored at -80°C.

**Bronchoalveolar lavage and tissue extraction**

Bronchoalveolar lavage (BAL) was performed with 800 µl chilled sterile PBS containing protease inhibitor (complete mini; MERCK). BAL fluid (BALF) and BAL cells were separated via centrifugation (500 RCF, 10 minutes, 4 °C). BALF supernatant was stored at -80 °C until further analysis and cell pellets were resuspended for leukocyte quantification. Lungs were flushed with 0.9% saline and left lung was harvested and stored at -80 °C for qPCR and protein quantification. Right lung lobes were divided into two parts, homogenized and used for bacterial load determination and leukocyte differentiation and quantification. Spleen and liver were also homogenized for determination of bacterial load.

**Lung permeability**

To determine lung permeability, total protein concentration was measured in BALF using the DC protein assay kit (BIO-RAD).

**Flow cytometry analysis**

Flow cytometry was used to differentiate leukocytes in BAL and lungs leukocytes as previously described (3). Briefly, lungs were flushed, and half of the right lung lobes was digested with DNAse (AppliChem GmbH) and collagenase (MERCK) and homogenized to prepare single cell suspension. After blocking for unspecific bindings (anti CD16/32, BD Biosciences), cells were stained with anti-CD11c-APC (N418, BD

Biosciences), anti-CD11b-PE-Cy7 (M1/70, BD Biosciences), anti-CD45-FITC (30-F11, BD Biosciences), anti-Ly6G-PerCP Cy5.5 (1A8, BD Biosciences), anti-Ly6C-BV510 (HK1.4, BioLegend), anti-MHCII-AF700 (M6/114.15.2, eBioscience), and anti-SiglecF-BV421 (1RNM44N, eBioscience). Neutrophils (PMN, polymorphonuclear leukocytes) were identified as $CD45^+$, $CD11c^-$, $CD11b^+$ and Ly-$6G^+$ cells, inflammatory monocytes as $CD45^+$, $CD11c^-$, $Ly^-6G^-$ and Ly-$6C^{high}$ cells, and alveolar macrophages as $CD45^+$, $CD11c^+$, $F4/80^+$ and siglec-$F^+$ cells(3). Counting beads (CountBright beads, Thermo Fisher Scientific) were used to quantify leukocytes according to manufacturer's instructions. Cells were acquired and analyzed using a FACS Canto II (BD Bioscience). Data were analyzed with FlowJo Version 10.1 software (BD Bioscience).

#### Cytokine and chemokine quantification

Cytokines were quantified in BALF and plasma using the multiplex bead-based immunoassay technique (LEGENDplex, BioLegend), according to manufacturer's instructions. Analysis was performed using a FACS Canto II (BD Bioscience) and data were analyzed with the LEGENDplex data analysis software (BioLegend).

#### Bacterial load determination

Serial dilutions of blood, BAL, and tissue homogenates (lung, liver and spleen) were performed, and samples were plated on Columbia Blood Agar plates with 5% sheep blood (BD Bioscience). Plates were incubated at 37° C under 5% $CO_2$ overnight. CFUs were counted on the following day. For bacterial load determination in liver and spleen, whole organs were used.

#### Histopathology

Duplicate experiments were performed for histological analyses of the lungs. Mice were anesthetized (160 mg/kg ketamine and 75 mg/kg xylazine), heparinized, and exsanguinated 48 h after infection. Tracheal ligation was performed to avoid alveolar collapse, and lungs were carefully harvested and fixed in 4% paraformaldehyde solution (pH 7.0). After fixation, lungs were routinely embedded in paraffin and cut into 5-µm-thick sections followed by dewaxing, dehydration and hematoxylin and eosin (H&E) staining. The degree of edema formation was assessed semi-quantitatively (0 = no edema, 1 = minimal edema, 2 = mild edema, 3 = moderate edema, 4 = severe edema). Three evenly distributed sections per lung were microscopically evaluated to assess edema formation. Histopathologic examination was performed by a European College of Veterinary Pathologists (ECVP) board-certified pathologist blinded to the study groups.

**Femoral bone marrow isolation**

For bone marrow (BM) isolation, 1 femur and tibia per animal were flushed, material was filtered, and cells were centrifuged with 1500 RCF at room temperature (RT). For PMN enrichment, positive selection was performed with the EasySep mouse neutrophil enrichment kit (Stem Cell Technologies) according to the manufacturer's instructions.

**Reactive oxygen species (ROS) quantification**

PMN ROS production was assessed with a luminol kinetic assay(4). Briefly, $5 \times 10^4$ isolated PMNs were plated in round clear-bottom 96-well plates, stimulated with PMA (100 nM; Thermo Fisher Scientific) and luminescence was recorded over 30 min in a SpectraMax L reader (Molecular Devices).

***Ex vivo* human lung tissue**

Fresh lung explants were obtained from patients undergoing lung resection at local thoracic surgery clinics. Written informed consent was obtained from all patients and the study was approved by the ethics committee at Charité - Universitätsmedizin Berlin, Germany (EA2/079/13). For infection with *S. pneumoniae*, tumor-free lung tissue was cut into small pieces (each ca. 0.1 - 0.2 g), weighed and incubated in RPMI 1640 medium at 37 °C with 5% CO2. Specimens were incubated for 24 h in RPMI 1640 with 10% heat-inactivated fetal calf serum. Subsequently, lung organ cultures were inoculated with 300 µl of prepared control or S.pn (1x106 CFU/ml) containing medium per 100 mg tissue, thereby assuring thorough stimulation of the tissue. The lungs were processed for further analysis 24 h after infection.

***In vitro* human alveolar epithelial cell model**

Human alveolar epithelial cells (A549 cells, ATCC CCL-185) were grown in RPMI medium (2 mM glutamine) supplemented with fetal calf serum (FCS). After cells reached confluence, media was replaced with plain media (no FCS) and cells were infected with live D39 wild-type (*S.pn* WT) or D39 *S.pn* pneumolysin-deficient (*S.pn* ΔPly) *S.pn* at MOI 50 for 4 hours. In separate experiments, A549 were challenged with pneumolysin, (PLY; 100 ng/ml) for 4 hours. Detailed procedures were described previously (5).

**Immunoblotting**

Proteins in cell lysates were separated by SDS-electrophoresis using 4-20% precast polyacrylamide gels (Genscript, NJ). Proteins were transferred to PVDF membranes and incubated with antibodies against IGF1R (Cell Signaling Tech) or β-actin (Sigma-Aldrich). Proteins were detected after incubation with HRP-conjugated secondary antibodies (Santa Cruz Biotechnology) and the ECL Western Blotting Substrate (Thermo Fischer Scientific). Densitometric analysis was performed using Image J.

**Gene expression analysis**

Flushed left lungs were homogenized with Gentle MACS M tubes (Miltenyi Biotec) in TRIzol and total RNA was extracted using an RNA purification kit (Zymo Research) according to manufacturer's instructions. cDNA was synthesized using the High-Capacity cDNA Reverse Transcription Kit (Applied Biosystems). qPCR was performed using the SYBR Green Master mix (Thermo Fisher Scientific) on a CFX96 real-time PCR detection system (BIO-RAD). GAPDH was used as housekeeping gene. The ΔΔct method was used and the fold change expression compared to PBS infected *Igf1r*$^{fl/fl}$ controls was calculated via 2^-ΔΔct. Predesigned primers were purchased from Qiagen (QuantiTect Primer Assays).

**Patient serum**

CAPnetz provided serum samples of patients with CAP where *S.pn* or influenza virus was identified. The study has been approved by the ethics committee of the Hannover Medical School (registration number 301-2008) and by the local responsible ethics committees of all study centers. All study participants suffered from confirmed CAP and were at least 18 years of age. Written informed consent was obtained from patients or their legal representatives. CAP was defined as working diagnosis of CAP provided by the enrolling physician, pulmonary infiltrate detected by chest X-ray, and at least two of the following five symptoms: 1) fever, 2) cough, 3) purulent sputum, 4) shortness of breath or need for respiratory support, or 5) crackling or rales on auscultation, dullness to percussion, or bronchial breathing. Control samples were obtained from healthy volunteers. Written informed consent was obtained from each patient, or each patient's legal representative. Serum levels of IGF1 and soluble IGF1R were quantified via respective ELISA kits (ab100545, ab100546, Abcam).

**Statistical analysis**

All experimental data are expressed as mean ± SEM. For comparison between two groups, Mann-Whitney U test was applied. For multiple comparisons, two-way ANOVA with Tukey's multiple comparisons test or one-way ANOVA with Tukey's multiple comparisons test was used depending on the variables. p values $< 0.05$ were considered statistically significant with *$p < 0.05$, **$p < 0.01$, ***$p < 0.001$. Analyses were performed using Prism 9.00 (GraphPad Software, USA).
